# Tau seeding in neurons enabled by transient endolysosomal perforations are confined within endolysosomes

**DOI:** 10.1101/2025.11.02.686178

**Authors:** Anwesha Sanyal, Gustavo Scanavachi, Elliott Somerville, Beren Aylan, Jose Inacio Costa-Filho, Forest Brooks, John R. Dickson, Bradley T. Hyman, Tom Kirchhausen

## Abstract

Pathogenic tau assemblies propagate by templated seeding. For endocytosed fibrils to initiate aggregation of endogenous tau, a breach must occur in the limiting membrane of an endosome or lysosome. To study the route by which internalized tau seeds access cytosolic monomers and to identify the site of aggregate growth, we imaged live human iPSC-derived neurons (iNs) expressing tau P301L–eGFP after exposure to recombinant tau pre-formed fibrils (PFFs) or Alzheimer’s disease (AD) brain–derived oligomers or fibrils. We detected seeded tau P301L– eGFP aggregation within late endosomes/lysosomes of iNs but not in undifferentiated iPSCs. Colocalization with a Dextran pH biosensor showed that the aggregates remained within the lumen of an intact, low-pH compartment. Reporters of endolysosomal injury and repair (endolysosomal recruitment cytosolic galectin-3 and the ESCRT-III component IST1) did not change during seeding. Volume focused-ion-beam scanning electron microscopy showed fibrillar material exclusively inside membrane-bounded endolysosomes, with no membrane discontinuities in the fibril-containing compartments and with no evidence of cytosolic aggregates. Because tau and α-synuclein can cross-seed, we adapted a HaloTag pulse–chase assay to test for the persistence of trans-membrane access. AD fiber–containing endolysosomes progressively recruited cytosolic α-synuclein–Halo over days, with heterogeneous incorporation histories consistent with recurrent, self-limited access events rather than persistent rupture or terminal sealing. Pharmacologic inhibition of the endolysosomal lipid kinase PIKfyve with apilimod suppressed seeded tau aggregation and prevented neuronal toxicity. These data indicate that templated conversion proceeds within acidic, membrane-intact endolysosomes; tau seeding in neurons is enabled by transient, self-limited endolysosomal perforations yet remains confined to the endolysosomal lumen, and it requires PIKfyve-dependent PI(3,5)P₂.

## INTRODUCTION

Tau assemblies propagate by a seeded, prion-like mechanism in cells and *in vivo*, in which internalized seed competent tau fibrils convert soluble cytosolic tau into cytosolic aggregates that ultimately spread across neural circuits (e.g^1–7^ Key unresolved questions concern the subcellular site and the conditions of templating. Work in cell systems has variously implicated endocytic uptake via heparan-sulfate proteoglycans, processing within the endolysosomal pathway, and seeding after vesicle injury or escape to the cytosol^4^.Related studies in glia reported lysosomal neutralization and nanoscale membrane injury after exposure to tau fibrils, promoting the view that lysosomal rupture may permit cytosolic access ^8^.

We recently showed that human iPSC-derived neurons (iNs)contain a small subset of endolysosomes with transient nanopores in the limiting membrane ^9^. These openings persist for ≥10 min and permit episodic access of cytosolic α-synuclein (α-syn) to the lumen, where internalized pre-formed fibrils (PFFs) of α-syn template further fibril growth. The newly extended fibrils are toxic, but they remain endolysosomal. Inhibition of the endosomal phosphoinositide kinase PIKfyve (e.g., by apilimod) blocks seeded aggregation and mitigates toxicity ^9^. These findings suggest a general route for lumenal seeding by internalized assemblies.

To test an analogous mechanism for tau, we asked whether iNs also support tau seeding by internalized tau assemblies and, if so, where does templated growth occur and how does it relate to endolysosomal openings. The results with tau parallel those with α-synuclein and lead to a coherent picture. Assemblies seeded by recombinant tau PFFs and AD brain–derived tau oligomers or fibrils remain within endolysosomes, which are acidic and membrane-intact, with no detectable rise in steady-state damage or markers of repair. These observations reconcile earlier models by locating templated growth inside endolysosomes and indicating that transient, reversible openings suffice to admit cytosolic monomers while preserving compartmental integrity.

To probe the dynamics of access, we adapted an optical pulse–chase strategy previously used to monitor nuclear pore biogenesis during mitosis ^10^ and applied it to endolysosomal seeding in iNs. Adding a membrane-permeable Halo ligand to cells expressing α-syn–HaloTag, followed by sequential incubations with spectrally distinct HaloTag ligands, we tracked incorporation into AD fibril–containing endolysosomes over days. The heterogeneous, lumen-confined recruitment histories are consistent with infrequent, self-limited, opening–closure cycles of the endolysosomal limiting membrane rather than persistent rupture or terminal sealing.

Finally, we showed that apilimod, an inhibitor of PIKfyve that lowers endolysosomal PI(3,5)P₂ and perturbs endolysosomal trafficking and ion homeostasis, suppresses tau seeding and rescues neuronal viability, acting upstream to limit templated conversion rather than downstream to enhance clearance. Prior work links PIKfyve modulation to diminished K18– PFF–driven aggregation, consistent with our findings^11^.

Our results localize seeded tau growth to closure-competent endolysosomes and identify endolysosomal lipid signaling as a point of potential intervention to limit aggregate templating.

## RESULTS

### Internalized Tau PFFs and AD brain–derived oligomers and fibrils seed tau in neuronal endolysosomes

We generated iPSCs and iPSC-derived neurons (iNs) stably expressing full-length tau P301L– eGFP, a fluorescent chimera fusion that enabled live-cell visualization of templated conversion of cellular monomeric tau into fibers, mediated by internalized tau PFFs ^7^. In both iPSCs and iNs, tau P301L–eGFP localized along microtubules, as expected for tau ^12–14^ (Fig. 1A, F).

**Figure 1.**
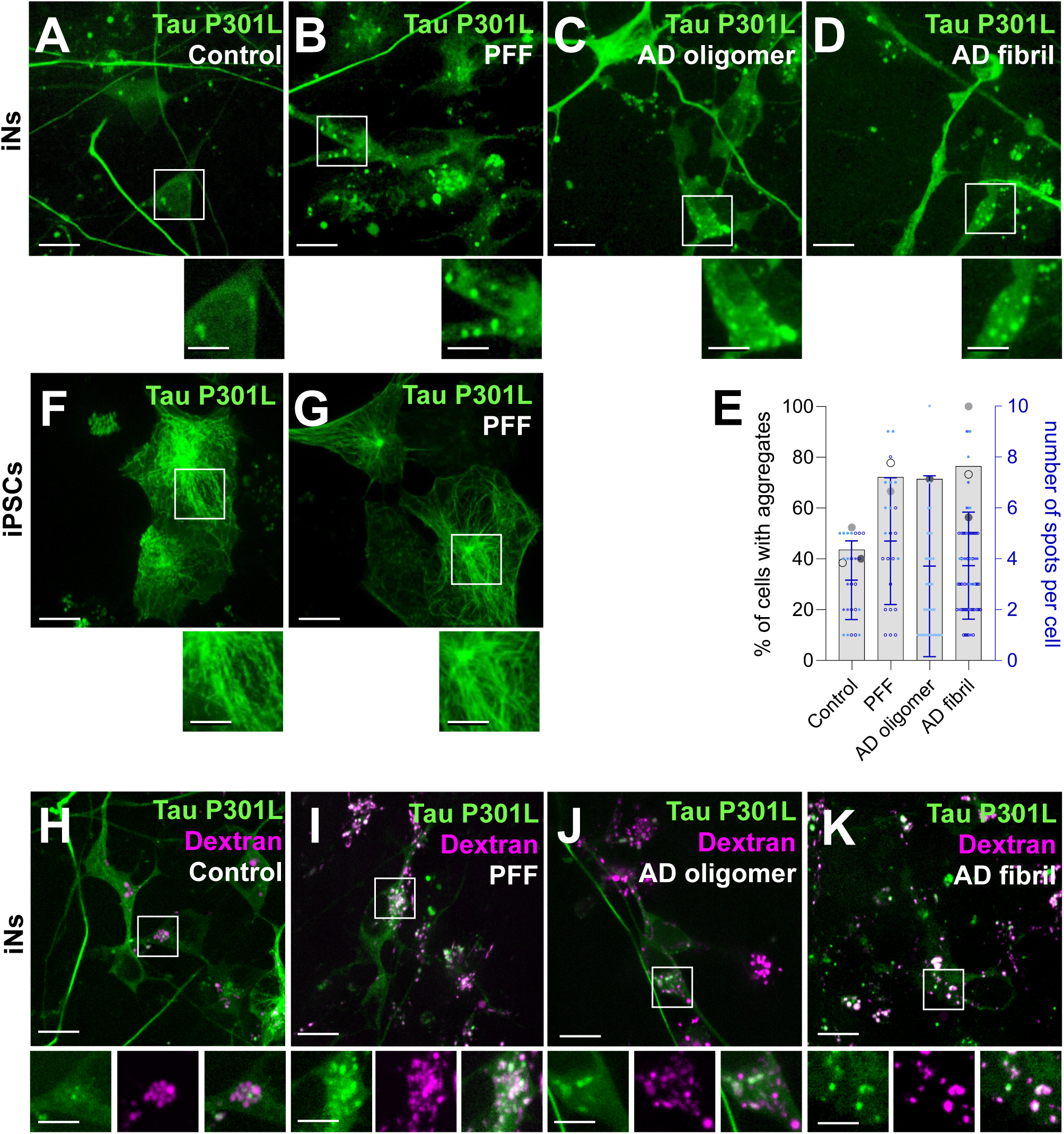
Tau P301L–eGFP aggregates form in endolysosomes of iNs exposed to seed competent recombinant tau PFFs or AD brain–derived oligomers or fibrils. **(A–D)** iNs expressing Tau P301L–eGFP show diffuse cytosolic fluorescence at baseline **(A)** and punctate aggregates after 1 day with 2 µg/ml PFFs **(B)** or 5 day with AD brain–derived oligomers **(C)** or fibrils **(D)**. Live-cell spinning-disk confocal images are maximum-intensity z-projections; images are representative of two biological replicates. Scale bar, 10 µm. Insets, 2×. **(E)** Quantification of the fraction of cells with aggregates; each dot is a biological replicate (48– 70 cells per condition), bars indicate mean. **(F, G)** iPSCs stably expressing Tau P301L–eGFP display diffuse signal in the absence **(F)** or after 1 day with 2 µg/ml PFFs **(G)**. Images are maximum-intensity z-projections; representative of one biological replicate. Scale bar, 10 µm. Insets, 2×. **(H–K)** iNs as in **A–D**, following a 2-h pulse with Alexa Fluor 647–dextran (20 µg/ml) immediately before imaging to mark the endolysosomal lumen: untreated **(H)**, 1 day with 2 µg/ml PFFs **(I)**, 5 day with AD oligomers **(J)**, or 5 day with AD fibrils **(K)**. Images are representative of two biological replicates (50–70 cells per condition). Scale bar, 10 µm. Insets, 2×.

Incubating iNs with recombinant tau PFFs ^15^ for 1 day or with AD brain–derived tau high molecular weight (HMW) oligomers or sarkosyl-insoluble tau fibrils ^16,17^ for 5 days and imaging with live-cell, 3D, spinning disc confocal microscopy revealed formation of tau-positive fluorescent puncta (Fig. 1B-D), which were absent in control cells not exposed to PFFs or AD-derived preparations (Fig. 1A, control). The puncta represented seeded tau aggregation. Under the same conditions, parental undifferentiated iPSCs showed no tau aggregation (Fig. 1F, G).

Thus, fibril-induced tau aggregation in iNs but not in iPSCs agreed with similar previous results observed in mouse hippocampal neurons expressing tau P301S-CFP and tau P301S-YFP treated with recombinant tau PFFs ^4^.

To define the subcellular site of the aggregates, we imaged live iNs expressing tauP301L–eGFP after a 2-h pulse with 6h chase with Alexa Fluor 647–dextran (Dextran-AF647), which at that time labels late endosomes and lysosomes ^9^ . In cells exposed to tau fibrils (Fig. 1I–K), but not in controls (Fig. 1H), most fibril-induced tau puncta colocalized with Dextran-AF647–positive vesicles. This held for both PFFs and AD oligomers or fibrils, indicating endolysosomal localization of the seeded aggregates as previously reported ^8^. As a critical control, adjacent iNs lacking tauP301L–eGFP showed no seeded fluorescent tau puncta (Fig. 1H), excluding the possibility that these spots arose from exogenous fluorescent tau released into the medium.

### Internalized tau fibrils do not compromise neuronal endolysosomes

We used two complementary assays to test whether internalized tau PFFs and fibrils perturb endolysosomal integrity. Using membrane damage and repair reporters in the first assay, we monitored recruitment of cytosolic Galectin-3 (Gal-3) (Fig, 2 A-E) and the ESCRT-III component IST1 (Fig. 2F-J) to the limiting membrane of endolysosomes ^8,18^. In iNs expressing Gal-3–eGFP or IST1–eGFP, both reporters were diffuse in the cytosol at baseline together with discrete puncta reflecting transient endolysosomal damage (Fig. 2A-D and 2F-I). Consistent with our previous work ^19^, untreated iNs also contained a small population of Gal-3–eGFP puncta (Fig. 2A). A 24-h incubation with PFFs or 5-day incubation with AD oligomers or fibrils did not increase the number of iNs with Gal-3–eGFP puncta or the number of Gal-3-eGFP spots per cell relative to the control (Fig. 2E). Although Ist1–eGFP puncta were significantly more abundant than Gal-3-eGFP puncta at baseline (Fig. 2F), their number did not change after comparable incubations with PFF, AD oligomers or AD fibrils (Fig. 2G–J). Thus, under our conditions, preformed tau assemblies internalized by iNs did not elicit a detectable increase in steady state endolysosomal membrane damage or repair as assessed by cytosolic Gal-3 access to luminal glycoconjugates or recruitment of Ist1, respectively.

**Figure 2.**
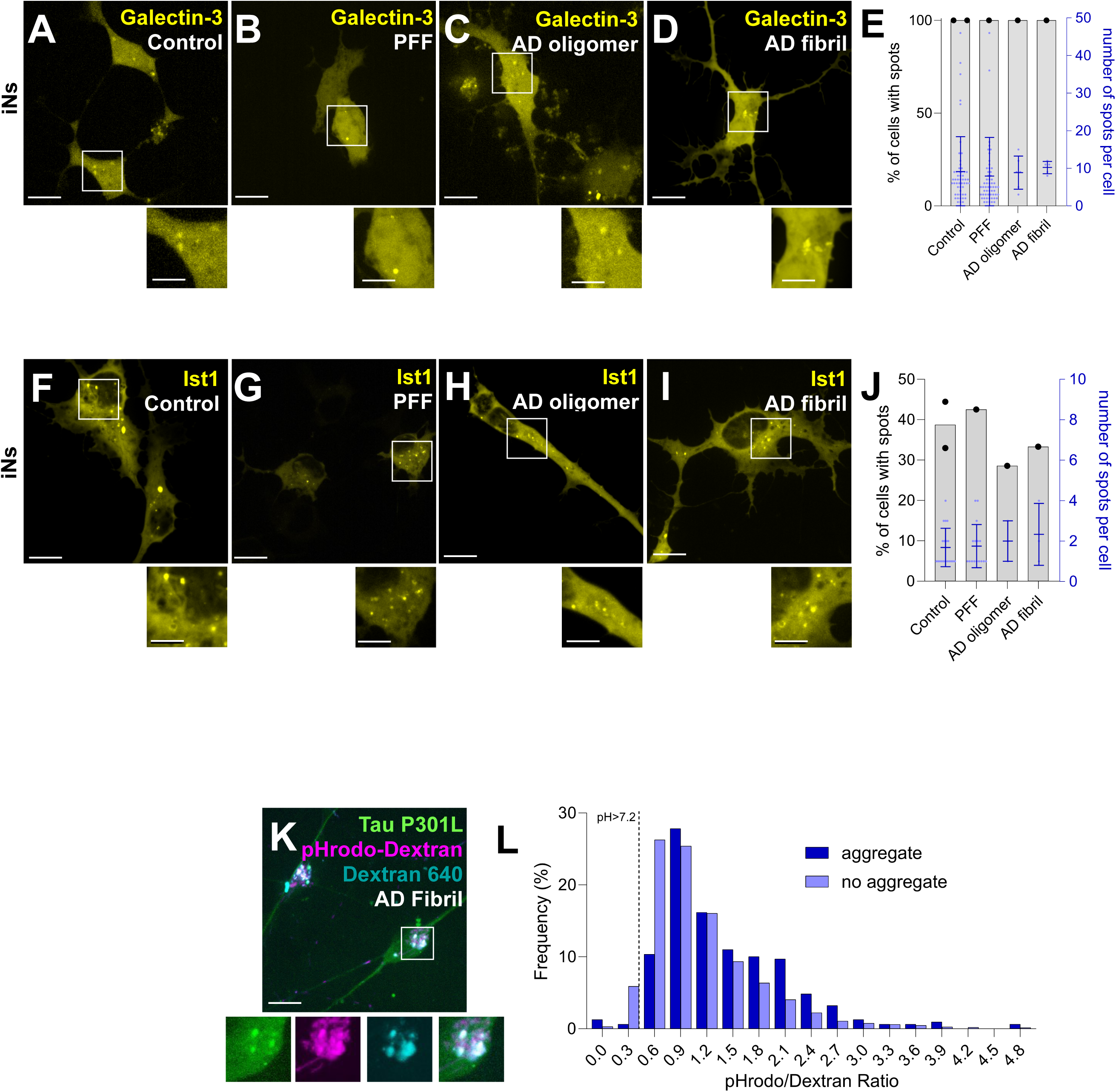
Seeded Tau P301L-eGFP aggregates locate to endolysosomes not actively damaged by tau oligomers or fibrils. **(A-E)** Live imaging of iNs expressing ESCRT III subunit, Ist1-eGFP incubated without **(A)** or **(B)** 2 µg/ml PFFs or **(C)** with AD oligomers or **(D)** fibrils for 1 day. Scale bar: 10 µm. Insets shown at 2× magnification. **(F–J)** Live imaging of iNs expressing eGFP-galectin-3 incubated without (F) or with (G) 2 µg/ml PFFs, (H) AD oligomers or (I) fibrils for 1 day. Scale bar: 10 µm. Insets show 2× magnification. The bar graphs in E and J quantifies the fraction of cells containing Ist1-eGFP or galectin-3- eGFP spots; each dot represents a biological replicate and the number of spots per cell; each dot represents a cell. 5-50 cells were analyzed. **(K**) iNs expressing Tau P301L-eGFP, incubated 5 days with AD brain derived fibrils ending with a a 2-h incubation with 20 µg/ml pH-sensitive pHrodo-Red Dextran and pH insensitive Dextran Alexa Fluor 647 before live cell imaging. Images are representative of 1 biological replicate with 87 cells. Scale bar: 10 µm. Insets show 2× magnification. **(L)** Bar graph quantifies distribution of acidic and neutral endolysosomes either containing or not Tau aggregates. 2595 spots from 87 cells were analyzed where 12% of the dextran colocalized with Tau aggregates.

For the luminal endolysosomal pH assay, we first incubated iNs with AD fibrils for 5 days, sufficient for seeded aggregate formation, then pulsed for 2 h with a a 6h chase immediately before live-cell imaging with a mixture of pH-sensitive dextran (Dextran-pHrodo) and a pH-insensitive dextran control (Dextran-640) (Fig. 2K). This ratiometric readout reports late endosome/lysosome luminal acidity ^19^. In untreated iNs, a very small number of all endolysosomes had neutral luminal pH reflecting the constitutively compromised endolysosomes (pH >7.2, Fig. 2L) we have reported previously, with minimal changes 5 days after exposure to AD-derived fibrils (Fig. 2L). Most (e.g. > ∼ 98%) seeded tau aggregates colocalized with acidic (pHrodo-positive) endolysosomes indicating fully enclosed endolysosomes (Fig. 2 K, L).

Thus, under our assay conditions, iNs incubated with tau PFFs, and AD oligomers or fibrils generated aggregates that remained confined within acidic, membrane-intact endolysosomes, with no detectable increase in constitutive endolysosomal damage (Gal-3) or repair (Ist1).

### 3D FIB-SEM shows internalized PFFs and seeded tau fibers remain confined to membrane-intact endolysosomes

We used 3D focused-ion-beam milling - scanning electron microscopy (FIB-SEM) to visualize random volumes from the soma of two iNs, imaged at 5×5×5 nm or 10×10×10 nm per voxel after high-pressure freezing and freeze substitution, to test whether internalized PFFs and their seeded fibers generated after one day of incubation remain confined to membrane-intact endolysosomes or also appear in the cytosol. Negative-stain EM verified that the PFF preparation used for the experiment appeared as clusters of linear filaments, each cluster comprising a variable number of filaments with lengths and cross-sectional widths of ∼ 10-20 nm, consistent with prior observations of similarly prepared recombinant tau PFFs ^20^.

In iNs, we found fibers made of clusters of linear fibrils exclusively within membrane-bounded compartments containing numerous 50-90 nm in diameter intraluminal vesicles, 50-90 nm in diameter, together with irregular membranous profiles characteristic of multivesicular bodies and endolysosomes ^21^ . In some of these structures, fibril ends abutted the limiting membrane, suggesting outward “pushing,” yet we found no discontinuities indicative of membrane rupture. We inspected the soma of 2 iNs with Neuroglancer -- their total volume of 1056x10^^3^ µm³ contained 17 endolysosomes with intraluminal fibrils, respectively, and 27 endolysosomes lacking detectable fibrils. To aid inspection, we segmented fibers with Labkit ^22^ and ASEM ^23^ and examined them manually; fibrils appeared as aligned clusters. Finally, despite robust identification of fibrils inside endolysosomal structures, we did not detect fibrils in the cytosol in any surveyed volume. These high-resolution volumetric data are consistent with our lower-resolution fluorescence microscopy, which placed seeded tau aggregates within intact endolysosomes and not in the cytosol.

### Recurrent endolysosomal damage–repair permits AD fibril–mediated seeding of cytosolic α-synuclein

We took advantage of our α-synuclein–Halo construct and the published observation that tau monomers can cross-seed α-syn fibrils ^24^ to test the reciprocal reaction: whether internalized AD fibrils could cross-seed recruitment of cytosolic α-syn–Halo. In iNs exposed to AD fibrils for 5 days, cytosolic α-syn–Halo was robustly recruited into endolysosomes, comparable to tau– eGFP recruitment under identical conditions (Fig. 4A).

We then used a dual-fluorescent HaloTag pulse–chase to report endolysosomal access events that admit cytosolic α-syn–Halo (“opening”) and their subsequent repair ("closing"). At t₀, we added tau PFFs and delivered a brief pulse of membrane-permeable HaloTag ligand 1 (HT1), which labeled at baseline cytosolic α-syn–Halo (and α-syn–Halo in the lumen of any accessible organelle). At the first detection of aggregates (t_agg_, about 4 days after fibril exposure), we gave a second HT1 pulse, labeling α-syn–Halo synthesized between t₀ and t_agg_, including any α-syn– Halo that had entered endolysosomes to form initial aggregates. At t_end_ (day 6), we pulsed with a spectrally distinct HaloTag ligand (HT2) to label α-syn–Halo synthesized after t_agg_, both in the cytosol and and in endolysosomal aggregates. HT1-only aggregates would indicate no post-t_agg_ access; mixed HT1/HT2 at a ratio exceeding the cytosolic ratio at t_end_ would indicate recurrent access both before and after t_agg_; HT1/HT2 equal to or less than the cytosolic ratio at t_end_ would indicate seeding after t_agg ._

For the first experiment, iNs expressing α-syn–Halo were continuously incubated for 7 days with AD brain–derived fibrils and pulsed for 10 min with Halo–JF657 on days 0 and 4 and with Halo– JF549 on day 6, followed by immediate live 3D imaging. Puncta consistent with endolysosomal localization appeared, with extensive colocalization of both Halo tags (Fig 5A). The fluorescence-intensity ratio histogram (Halo–JF657/Halo–JF549), referenced to the cytosolic ratio at day 6 (6.7 ± 3.7), showed a continuum: a fraction overlapping the cytosolic ratio, a dominant population with higher ratios (median ∼20), and a few puncta at 40–50; no puncta contained only the first dye (Fig 5B). We interpret puncta with cytosol-like ratios as aggregates seeded between days 4–6; higher ratios indicate seeding that had begun by ∼day 4 (when only Halo–JF657–labeled α-syn–Halo was available). From these data, we estimated the number of α-syn–Halo molecules incorporated per punctum based on the incorporation ratio of Halo– JF657 and Halo–JF549 (Figs. 5C, D): up to ∼3,500 molecules for Halo–JF657 over days 0–6 and up to ∼ 250 for Halo–JF549 over days 4–6. Although we cannot distinguish single versus multiple seeding bursts, the heterogeneous dye ratios imply sporadic endolysosomal access events; if incorporation proceeded at a similar monomer-addition rate throughout, a single continuous fiber would have an upper-bound length of ∼1,750 nm (3,500 molecules ÷ ∼2 molecules·nm⁻¹), whereas seeding of ∼6 fibers would yield extensions of ∼300 nm each, dimensions within the range of clustered fibrils observed by FIB-SEM in iNs seeded with tau PFFs (Fig. 3). For this analysis, we assumed ∼80% covalent labeling of the genetically encoded HaloTag in live cells by the Janelia Fluor HaloTag ligand, based on previously reported results for other HaloTag proteins^10,25–27^.

**Figure. 3 (Associated with Video 1).**
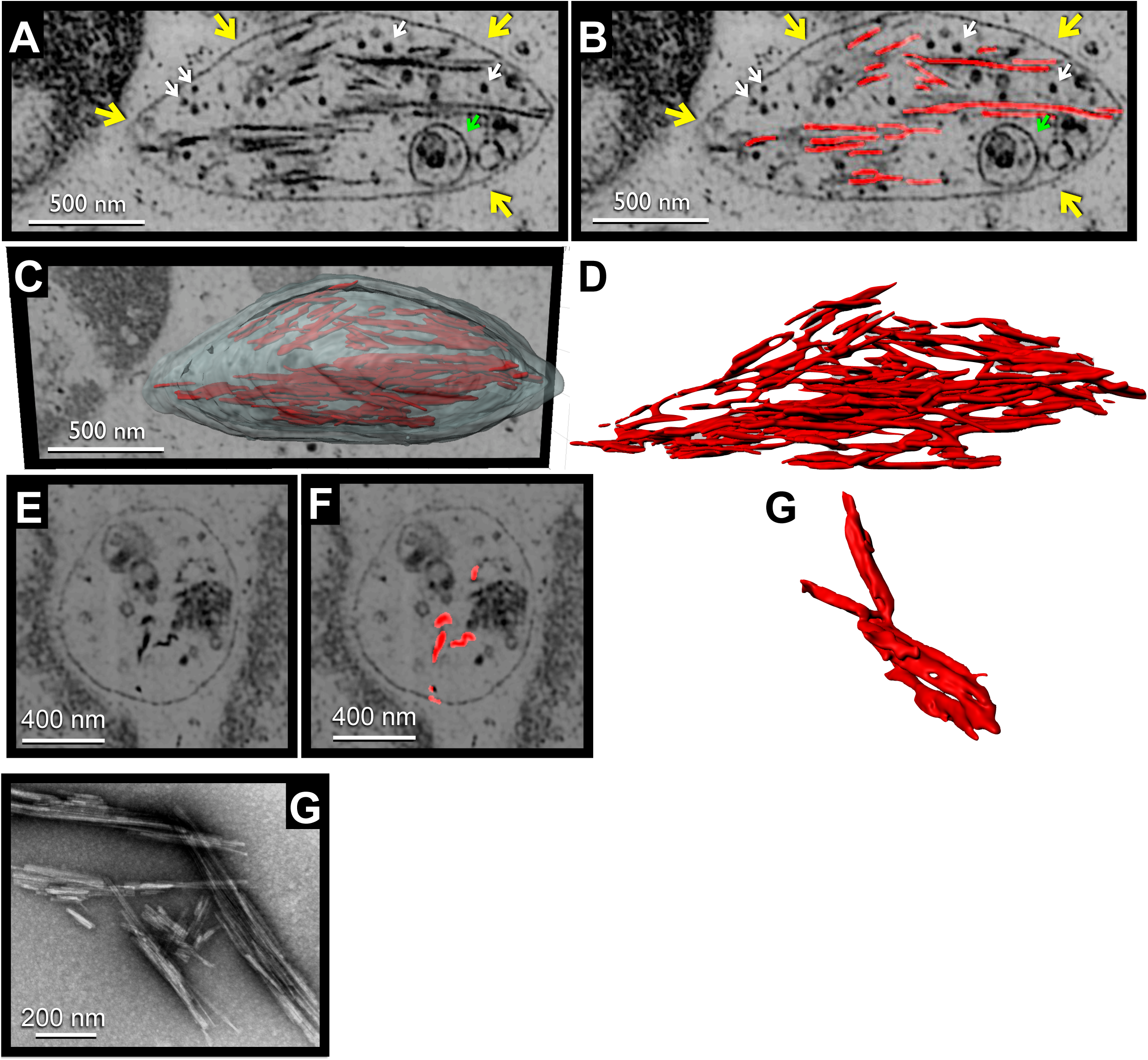
3D FIB-SEM of an endolysosome in an iN processed by high pressure freeze substitution containing internalized tau PFFs. iNs were incubated for 1 day with tau PFFs before imaging at 5 × 5 × 5 nm/voxel. **(A)** Representative denoised single-plane section through of an endolysosome bounded by an intact limiting membrane (yellow arrow), with intraluminal vesicles (white arrow), heterogeneous internal membranes (green arrow), and cross-sections of fiber/fibrils (red). Scale bar, 500 nm. **(B)** Same as **(A)** including single plane segmentation outline of the fibrils contained within the endolysosome generated with Labkit. Scale bar, 500 nm. **(C)** 3D surface rendering of fiber/fibrils superimposed on a single-plane section of the endolysosome. Scale bar, 500 nm. **(D)** 3D surface rendering of the segmented fibrils depicted in **(C)**. **(E,F)** Another example of and endolysosome containing tau fibrils: **(E)** without segmentation overlay; **(F)** with single-plane segmentation outlining the fibrils. Scale bar, 400 nm. **(G)** Representative negative-stain TEM of tau PFFs, stained with 1% uranyl acetate. Scale bar, 200 nm.

**Figure 4.**
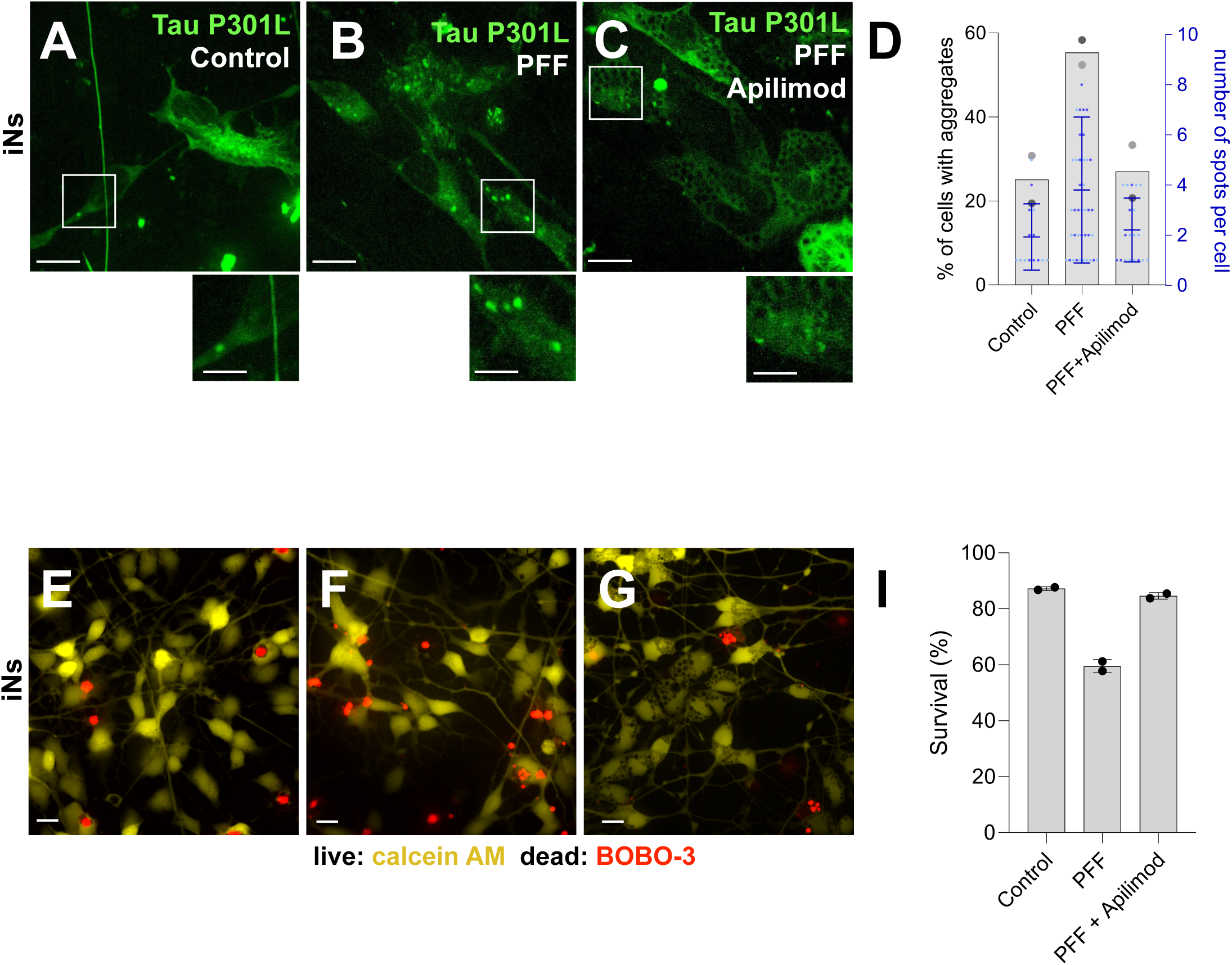
PIKfyve inhibition prevents PFF-induced Tau aggregation and cell death. **(A–C)** iNs expressing Tau P301L–eGFP were incubated for 1 day in the absence of PFFs **(A)**, with 2 µg/ml PFFs **(B)**, or with 2 µg/ml PFFs plus 1 µM apilimod **(C)**. Scale bar, 10 µm. Insets, 2×. **(D)** Plots for fraction of cells with Tau fluorescent spots indicative of aggregates; each dot denotes the average from a biological replicate (62–78 cells per condition). Number of eGFP fluorescent spots color coded for per each biological replicate is shown. **(E–G)** iNs were incubated for 2 day without PFFs **(E)** or with 2 µg/ml PFFs in the presence or absence of 1 µM apilimod **(F, G)**. Live cells were labeled with calcein AM; dead cells with BOBO-3 iodide. Images are representative of two biological replicates. Scale bar, 10 µm. **(H)** Quantification of proportion of live and dead iNs for the indicated treatments; data is from two biological replicates, 20 fields per condition (∼6,500 cells total).

**Figure 5.**
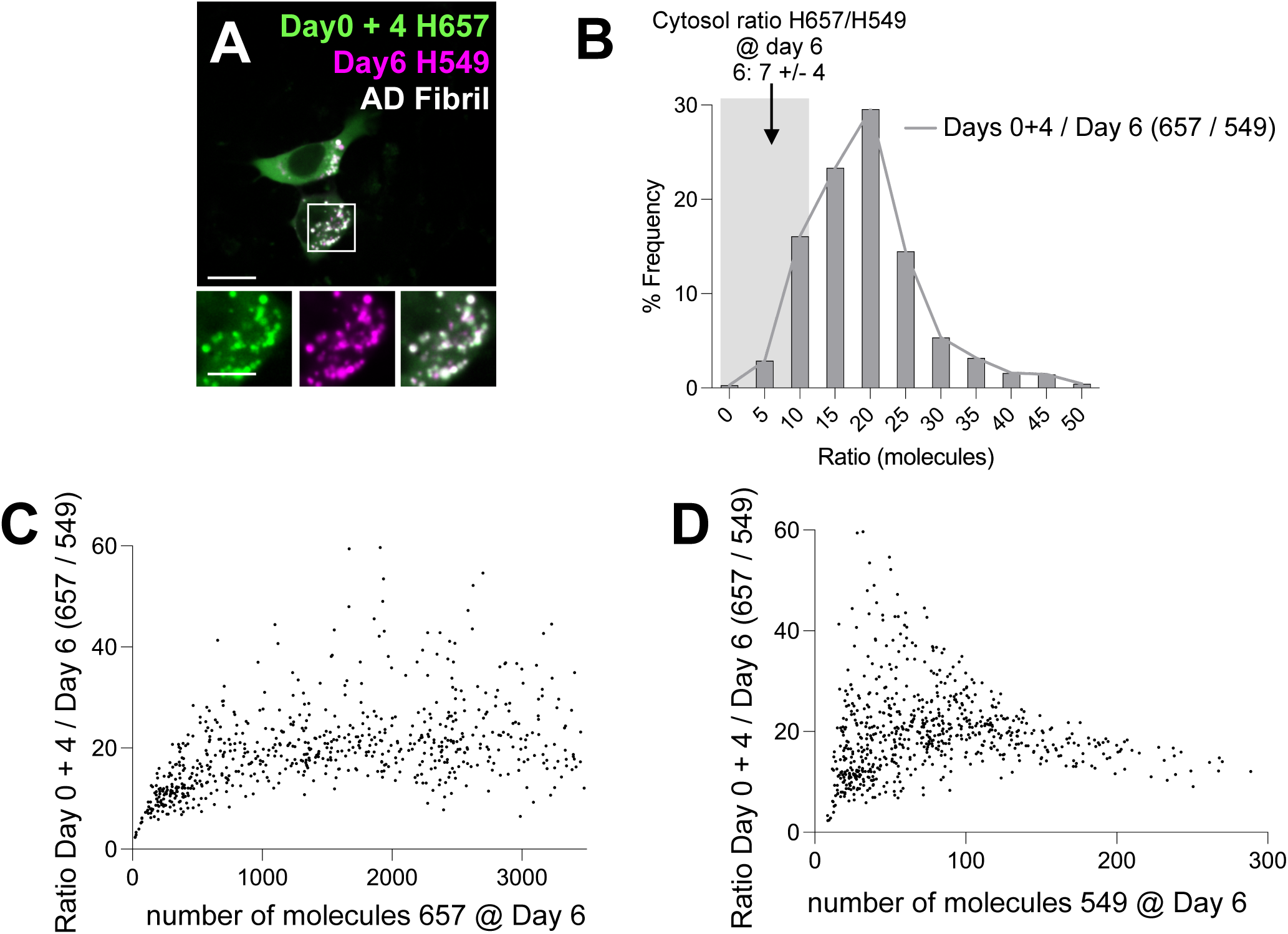
Recurrent endolysosomal damage–repair permits AD fibril–mediated seeding of cytosolic α-synuclein. **(A)** Live imaging of iNs expressing α-synuclein-Halo incubated for 7 day with AD brain fibril. α- synuclein-Halo was labeled with JF657 on day 0 and day 4 and with JF549 on day 6. Scale bar, 10 µm. Insets, 2×. **(B)** Frequency distribution of ratio of number of molecules of α-synuclein-Halo labeled with JF657 and JF549 in the tau aggregates imaged at day 6. The labelling range for the cytosol on day 6 is indicated by the gray shaded region. **(C,D)** XY scatter graphs showing correlation between ratio of number of molecules of JF657 and JF549 with **(C)** number of molecules of JF657 or **(D)** number of molecules of JF549 imaged on the day 6. Image and graphs are representative of data from the tau aggregates in one biological replicate with 45 cells.

For the second experiment, we added Halo–JF657, Halo–JF549, and Halo–JF503 for 10 min at days 0, 2, and 4, respectively, with imaging on days 2 and 4. As expected, puncta appeared at day 4 but not at day 2 (Fig. 6A). The outcome mirrored the first experiment: all puncta contained all three dyes in varying amounts, indicating incorporations at day 4 of up to ∼3,500, ∼1200, and ∼80 α-syn–Halo molecules labeled on days 0, 2, and 4, respectively, with a detection limit of ∼15 molecules per punctum (Fig 6C-H). These results are consistent with the first experiment. For this analysis, we assume ∼80% covalent labeling of the genetically encoded HaloTag in live cells by the Janelia Fluor HaloTag ligand, based on previously reported results for other HaloTag proteins ^10,25–27^.

**Figure 6.**
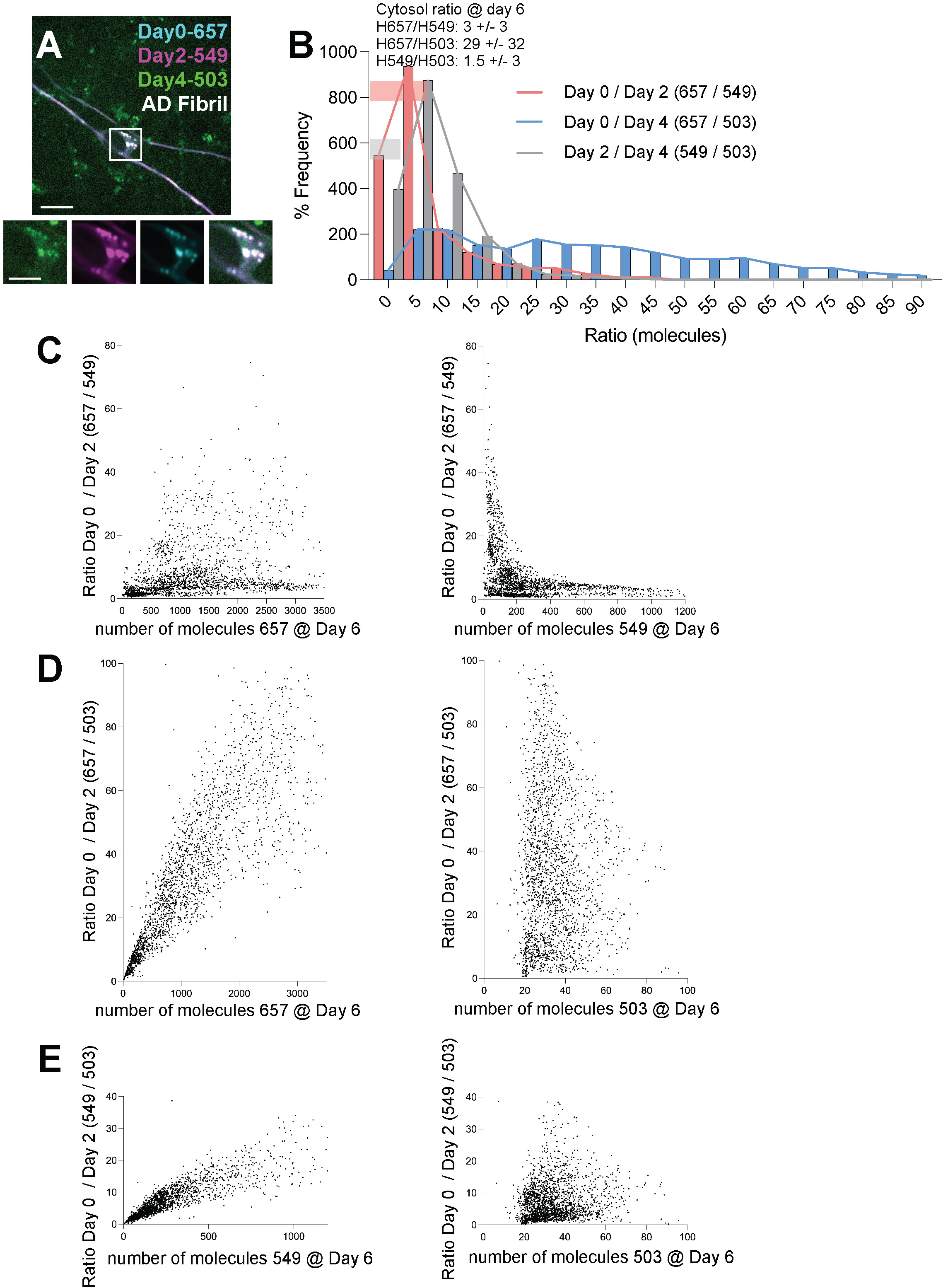
Recurrent endolysosomal damage–repair permits AD fibril–mediated seeding of cytosolic α-synuclein. **(A)** Live imaging of iNs expressing α-synuclein-Halo incubated for 5 day with AD brain fibril. α- synuclein-Halo was labeled with JF657 on day 0, JF549 on day 2 and with JF503 on day 4. Scale bar, 10 µm. Insets, 2×. **(B)** Frequency distribution of ratio of number of molecules of α-synuclein-Halo labeled with either JF657 and JF549, JF657 and JF503, or JF549 and JF503. **(C-E)** XY scatter graphs showing correlation of **(C)** ratio between number of molecules of JF657 and JF549 and (left) number of molecules of JF657 or (right) of JF549; of **(D)** ratio of number between molecules of JF657 and JF503 with (left) number of molecules of JF657 or (right) of JF503; and of **(E)** ratio between number of molecules of JF549 and JF503 and (left) number of molecules of JF549 or (right) of JF503 imaged on day 4. Image and graphs are representative of data from one biological replicate with 20 cells.

The continuous spread of dye ratios among puncta indicates that seeded recruitment of α-syn– Halo monomers into endolysosomes containing internalized AD fibrils is asynchronous across endolysosomes. The data also show that a seeded fiber or a cluster of fibers within an endolysosome can incorporate monomers for at least 6 days. During this interval, we did not observe terminally sealed compartments unable to incorporate cytosolic α-syn-Halo (e.g. puncta labelled only with the first Halo Tag). Considering the acidic pH of all endolysosomes containing seeded fibers (see above), the simplest interpretation is that endolysosomal membrane openings permitting cytosolic access are transient. These data do not exclude fusion events between endolysosomes over the course of the experiment.

### PIKfyve inhibition with apilimod prevents seeded tau aggregation and rescues viability

We asked whether inhibiting the endolysosomal lipid kinase PIKfyve in iNs could modulate tau seeding and toxicity in the same way it blocks α-syn seeding and toxicity ^9^. iNs expressing Tau P301L–eGFP were exposed to recombinant PFFs (1 day) to induce seeded aggregation in endolysosomes. PFF treatment led to many more cells with Tau P301L–eGFP puncta than in untreated controls (Fig. 4A, B, D), elevated puncta count per cell, and progressive cell death, with ∼50% BOBO-3–positive (nonviable) cells versus ∼10% in untreated controls, and a corresponding decrease in calcein-AM–positive cells (Fig. 4I). Co-incubation with 1 μM apilimod reduced the number of cells with tau P301L–eGFP puncta to values comparable to those for control iNs not incubated with PFFs and reduced PFF–induced cell death to control levels as well (Fig. 4C, D, I). Thus, pharmacological inhibition of PIKfyve with apilimod prevents formation of tau aggregates seeded by recombinant PFFs and protects iNs from tau seed– mediated neurotoxicity.

## DISCUSSION

Our data place the initiating steps of tau templating within the endolysosomal system of human iPSC-derived neurons (iNs). Internalized recombinant tau PFFs and human AD brain–derived assemblies seed tau in iNs but not in undifferentiated iPSCs, and the resulting aggregates remain confined to acidic, membrane-bounded endolysosomes as evidenced by live-cell fluorescence microscopy and by 3D FIB-SEM at nanometer precision. We did not detect cytosolic fibrils within the cytosol of the sampled volumes. These observations argue that endolysosomes are not merely transit stations but the principal sites of early neuronal tau templating after uptake of misfolded tau from the environment.

We infer that endolysosomal membranes in this setting undergo transient, rather than catastrophic, breach. Recruitment of galectin-3, a sensor of limiting-membrane damage, did not increase, indicating no rise in lumen glycan exposure. The number of IST1 puncta, an ESCRT-III component, remained unchanged, consistent with invariant ESCRT-III–mediated repair. The simplest model that explains endolysosomal templating of cytosolic tau is recurrent, self-limited openings that admit cytosolic tau monomers and then close, with prompt restoration of lumen acidity. This “damage–repair cycle” reconciles efficient templating with retention of organelle integrity and accounts for the persistence of seeded aggregates within endolysosomes.

The presence of tau P301L–eGFP within aggregates indicates that cytosolic tau accesses the endolysosomal lumen and undergoes templated misfolding upon encountering resident fibrils, driving elongation. This mechanism contrasts with models that suggest rapid escape of fibrillar seeds into the cytosol followed by growth there (e.g.,^28–31)^. Because mature fibrils are ultimately cytoplasmic, we infer that endolysosomes later become sufficiently compromised to release their contents, but not on the timescale of our experiments.

The α-syn–Halo pulse–chase assay tracks, in real time, the coupling between endolysosomal aggregation and the exposure of internalized fibrils to cytosolic α-syn monomers. Distinct HaloTag pulses mark monomers available before, between, and after defined timepoints; their unequal incorporation into endolysosomal puncta reveals asynchronous access events across compartments and sustained monomer addition over days. The absence of initial HaloTag-only puncta and the broad ratio distribution with respect to the spectrally distinct HaloTags added several days later, both support recurrent, transient openings rather than a single, irreversible rupture. Estimated incorporation numbers and fiber-length bounds fall within the size range of clustered fibrils we visualize by FIB-SEM, linking molecular accrual to ultrastructure.

The endolysosomal pH measurements, repair markers, and volumetric FIBSEM define a regime of templated assembly by recombinant tau PFFs and AD-patient derived oligomers and fibrils behind an intact yet intermittently accessible limiting endolysosomal membrane and suggests that neuron-intrinsic repair capacity can preserve acidity even during active templating. Our regime differs from reports of neutralized lysosomes by internalized recombinant tau PFFs ^8^, for which *in vitro* fibrillization, in contrast to ours, was aided by heparin.

Our results also suggest a mechanism for selective vulnerability: small shifts in membrane opening frequency, ESCRT-mediated repair efficiency, endolysosomal fusion dynamics or stimulation of autophagy of compromised endosomes could tip the balance between contained templating and cytosolic escape. Inhibiting PIKfyve with apilimod disrupts this templating regime by blocking endolysosomal tau seeding and restoring viability. Because PIKfyve regulates PI(3,5)P₂, endolysosomal trafficking, and lumenal ion homeostasis, these data implicate the endolysosomal lipid milieu as a tunable determinant of seeding competence. The concordance of aggregation blockade with neuroprotection argues that targeting endolysosomal lipid signaling can act upstream of fibril amplification, not solely downstream at aggregate clearance.

We propose a working model in which continued intraluminal fiber elongation converts transient, repairable openings into unresolvable lesions that trigger a pro-death signal. In related α- synuclein assays, cytosolic aggregates in the absence of endolysosomal aggregates are not detectably toxic in human iNs ^9^, suggesting that the lethality signal originates at damaged endolysosomes; whether tau follows the same rule remains to be determined. These results position endolysosomes as the site of tau templating and as the likely source of the injury signal.

In summary, we have found that neuronal tau templating occurs within acidic, membrane-intact endolysosomes that undergo one or more cycles of damage-repair. We propose that transient openings over a period of days admit cytosolic monomers while preserving organelle integrity. Perturbing endolysosomal lipid signaling by inhibiting PIKfyve disables this pathway and protects the neurons. Finally, we suggest that damage–repair cycles in endolysosomes are a general driver of seeded assembly in neurons and an accessible therapeutic target for interfering with proteopathic spread.

Limitations: We are aware that fluorescent tags may alter protein behavior; our assays measure access and incorporation above defined detection limits and cannot exclude rare cytosolic escape below those limits; the pulse–chase design infers opening frequency from dye ratios rather than direct visualization. Nonetheless, independent readouts such as endolysosomal pH, ESCRT-III recruitment, quantitative monomer accrual, and ultrastructural confinement converge on the same model.

## MATERIALS AND METHODS

### Plasmids

pRK5-EGFP–Tau(P301L) (Addgene 46908) and pEGFP–hGalectin-3 (Addgene 73080) were obtained from Addgene. α-syn-Halo was generated by replacing YFP with Halo in α-syn-YFP- pcDNA3.1 coding sequence using NEBuilder HiFi DNA Assembly (NEB, E5520) according to the manufacturer’s instructions.

### Inhibitors

Apilimod (MedChemExpress, HY-14644) and a protease/phosphatase inhibitor cocktail (Thermo Fisher Scientific, 78440) were used as indicated.

### Cells

Human BR33 iPSCs, described previously ^32,33^ , were a gift from Dr. Tracy Young-Pearse (Brigham and Women’s Hospital, Boston, MA, USA). We maintained iPSCs in StemFlex medium (Life Technologies, A33493). For plating, we coated tissue-culture ware with Growth Factor–Reduced Matrigel for iPSCs (Corning, 354320) or standard Matrigel for iNs (Corning, 354234) by resuspending 0.5 mg Matrigel in 5 ml cold DMEM/F12, filtering through a 40-µm strainer (Corning, 352340), and applying 6 ml to a 10-cm dish (final ≈8.7 µg/cm²).

For iN induction, iPSCs were plated at 1×10^5 cells/cm² and co-transduced with lentiviruses encoding pTet-O-NGN2-puro and FUdeltaGW-rtTA, as described by Zhang et al. (2013). After 2 days, cells were expanded to confluence in a 10-cm dish and cryopreserved.

For production runs, we thawed transduced iPSCs onto Matrigel-coated 10-cm plates at 2×10^6 cells/plate in StemFlex supplemented with 10 µM ROCK inhibitor (Stemcell Technologies, 72304). Day 1: medium was replaced with KnockOut medium (Gibco, 10829-018) containing KnockOut Serum Replacement (Invitrogen, 10928-028), 1% MEM non-essential amino acids (Thermo Fisher Scientific, 11140-050), 1% GlutaMAX (Thermo Fisher Scientific, 35050-061), 0.1% β-mercaptoethanol (Invitrogen, 21985-023), and 2 µg/ml doxycycline (Sigma-Aldrich, D9891) to induce NGN2. Day 2: medium was changed to 1:1 KSR:N2B (DMEM/F12 [Thermo Fisher Scientific, 11320-033] with 1% GlutaMAX, 3% dextrose, and N2-Supplement B [Stemcell Technologies, 07156]) supplemented with 5 µg/ml puromycin (Life Technologies, A11138-03) and 2 µg/ml doxycycline. Day 3: N2B plus B27 (Life Technologies, 17504-044; 1:100), 5 µg/ml puromycin, and 2 µg/ml doxycycline. Day 4: iNs were frozen in Neurobasal medium (Gibco, 21103-049) supplemented with B27 (1:50), BDNF (Peprotech, 450-02; 10 ng/ml), CNTF (Peprotech, 450-13; 10 ng/ml), GDNF (Peprotech, 450-10; 10 ng/ml), 10 µM ROCK inhibitor, 5 µg/ml puromycin, and 2 µg/ml doxycycline (hereafter “NBM”), with 10% DMSO. We performed experiments on iNs 14–21 day after thaw, maintaining them in NBM.

### Lentivirus preparation and transduction

We seeded HEK293T cells at 2.5×10^6 cells per 15-cm dish in DMEM (Thermo Fisher Scientific, 11965118) supplemented with 10% FBS (Atlanta Biologicals, S11150H), 1× GlutaMAX (Thermo Fisher Scientific, 35050061), 1× sodium pyruvate (Thermo Fisher Scientific, 11360070), and 1× MEM non-essential amino acids (Thermo Fisher Scientific, 11140050). The next day, we transfected cells with Lipofectamine 2000 (Thermo Fisher Scientific, 12566014) according to the manufacturer’s instructions.

For each 15-cm dish, we combined 24 µg of each lentiviral transfer plasmid, 18 µg psPAX2 (Addgene 12260), and 12 µg pMD2.G (Addgene 12259) with 135 µL Lipofectamine 2000 in 6.75 mL Opti-MEM (Thermo Fisher Scientific, 51985091). After a 20-min incubation at room temperature, we added the mixture dropwise to cells in 14 mL complete medium. Six hours later, we replaced the medium with 14 mL fresh complete medium and, 12 h thereafter, collected supernatants. We cleared debris by centrifugation (500×g, 5 min, 4 °C), aliquoted the clarified supernatant (1 mL), snap-froze in liquid nitrogen, and stored at −80 °C.

For transduction, we added 1 mL thawed viral supernatant to ∼8×10^5 iPSCs seeded 18 h earlier in 6-well plates in 1 mL mTeSR1 (Stemcell Technologies, 85850). Twelve hours post-infection, we replaced the medium with StemFlex (for iPSCs) and applied selection with either blasticidin (5 µg/mL) or puromycin (5 µg/mL), according to the vector, for 4 day. We then cultured surviving cells for 7 day without antibiotics and cryopreserved them in the same medium containing 10% DMSO.

### Tau PFFs, AD oligomers and AD fibrils

Seed competent tau fibrils were prepared from recombinant human tau 297-391 expressed in *E. coli* using buffer and shaking conditions as described ^15^. Sarkosyl insoluble AD fibers (classic paired helical filaments) were isolated from unfixed human Alzheimer brain and generated by the extraction protocols as previously described ^16,17^. Seed competent AD oligomeric tau was isolated from the same brains by size exclusion chromatography of human AD brain lysate^34^.

### Transfection of iNs

We thawed and plated 4×10^4 iNs per well in 8-well chambered coverslips (Cellvis, C8-1.5H-N). We transfected cells with ViaFect (Promega, E4981) according to the manufacturer’s instructions. For each well, we mixed 0.2 µg plasmid DNA with 0.6 µL ViaFect in 100 µL Opti-MEM (Thermo Fisher Scientific), incubated 20 min at room temperature, and added the mixture dropwise to cells in 200 µL NBM. After 24 h, we replaced the medium with 200 µL fresh NBM. Unless noted, experiments were performed 14–21 day after thaw.

### Immunofluorescence

Cells on coverslips were rinsed in 1× PBS and fixed in 4% (wt/vol) paraformaldehyde for 30 min at room temperature. We permeabilized with 0.1% (vol/vol) Triton X-100 in 1× PBS for 5 min, then blocked in 5% (vol/vol) BSA in 1× PBS for 30 min. Primary antibody were applied for 1 h at room temperature in 5% BSA/1× PBS: LC3 (1:200; Abcam ab48394), After three 10-min washes in 1× PBS, we incubated with Alexa Fluor–conjugated secondary antibodies for 1 h at room temperature (Thermo Fisher Scientific: goat anti-rabbit A11008), followed by three additional 10-min washes in 1× PBS. Samples were imaged immediately in 1× PBS on a spinning-disk confocal microscope or stored at 4 °C in 1× PBS and imaged the next day.

### In vivo pH imaging and pH calibration

For pH calibration ^19,35,36^ , we loaded iNs for 2 h with pHrodo Red–dextran and Alexa Fluor 647– dextran (20 µg/ml each), washed three times in 1× PBS, and equilibrated for 15 min at room temperature in universal buffer (10 mM HEPES, 10 mM MES, 10 mM sodium acetate, 140 mM KCl, 5 mM NaCl, 1 mM MgCl₂) containing 10 µM nigericin (Cayman 11437) and 10 µM monensin (Cayman 16488). Buffer was titrated to pH 7.2, and we acquired volumetric images from two biological replicates and computed the mean pHrodo/Alexa Fluor 647 fluorescence ratio. Ratiometric quantification followed the Image analysis section.

We estimated mean pHrodo/Alexa Fluor 647 fluorescence ratio from a single time point by loading iNs with pHrodo Red–dextran (20 µg/ml; pKa ≈ 6.8) and Alexa Fluor 647–dextran (20 µg/ml) for 2 h at 37 °C, followed by a 6 h chase. After three washes in 1× PBS, we transferred cells to phenol red–free Neurobasal medium supplemented with 1% B-27, 10 ng/ml BDNF, 10 ng/ml CNTF, and 10 ng/ml GDNF. We then acquired live-cell ratiometric, whole-volume images at 37 °C on a spinning-disk confocal microscope as outlined in image analysis section. The range of ratios with estimated pH > 7.2 was estimated from the single pH point calibration value.

### Fluorescence Pulse–Chase Labeling

4×10^4 αSynuclein-Halo expressing iNs were plated per well of an 8-chamber slides and leabled with 200nM of different Janelia Fluor (JF) dyes for 30minutes at 37°C, followed by 3 washes with media for 10minutes each, For the 5-day experiment, iNs were pulsed sequentially with JF657 (day 0), JF549 (day 3), and JF503 (day 5). Cells were washed three times maintained under standard conditions until live imaging on day 5. For the 7-day experiment, cells were pulsed with JF657 on days 0 and 4, and JF549 on day 6, following the same labeling and wash procedures. Live imaging was performed on day 6.

### Cell survival assay

We determined the proportion of live and dead iNs by plating ∼40,000 cells on Matrigel-coated 8-chamber glass slides (C8-1.5H-N; Cellvis) and culturing them for 10 days after differentiation, followed by incubation with or without Apilimod and PFF for up to 2 days as previously described ^9^ . At the end of this step, cells in 50 μl of imaging media (phenol red-free Neurobasal medium supplemented with 1% B-27, 10 ng/ml BDNF, 10 ng/ml CNTF, and 10 ng/ml GDNF) were incubated for 15 min at room temperature with 50 μl of 2x stock solution of the LIVE/DEAD Cell Imaging Kit 488/570 (R37601; Thermo Fisher Scientific) containing Calcein AM for live cell staining and BOBO-3 Iodide for dead cell staining. Following incubation, spinning-disc confocal imaging at 40× magnification was performed, and the live and dead cells were counted.

### Spinning-disc confocal imaging

We plated 4×10^4 cells per well in 8-well chambered coverslips (Cellvis, C8-1.5H-N) pre-coated with Matrigel for iNs or growth factor–reduced Matrigel for iPSCs. iNs were imaged in phenol red–free Neurobasal (Thermo Fisher Scientific, 12348017) supplemented with 1% B27, 10 ng/mL BDNF, 10 ng/mL CNTF, and 10 ng/mL GDNF while iPSCs were imaged in FluoroBrite medium containing 10% FBS and 25 mM HEPES (pH 7.4). Live-cell imaging was performed at 37 °C in a humidified, temperature-controlled chamber with 5% CO₂; fixed samples were imaged at room temperature.

Image acquisition was controlled with SlideBook 6.4 (Intelligent Imaging Innovations, 3i) using the following microscope configurations:

a. Live-cell: Marianas system built on a Zeiss Axio Observer Z1 microscope with an additional 1.2× magnification lens; 20×/0.5, 40×/0.75, and 63×/1.4 Apochromat objectives; a Yokogawa CSU-XI spinning-disk unit; a heated stage (OKO Lab); a 3i spherical aberration correction system; and a 3i LaserStack with diode lasers at 488 and 560 nm (150 mW each), and 640 nm (100 mW). Z-stacks were acquired at 270-nm steps with 10-100 ms exposures using a sCMOS camera (Teledyne Photometrics Prime 95B).
b. Fixed-cell: Marianas system built on a Zeiss Axio Invert 200 M microscope with an additional 2× magnification lens; a 63×/1.4 Plan-Apochromat objective; a Yokogawa CSU-22 spinning-disk unit; and solid-state lasers at 491, 561, and 660 nm. Z-stacks were acquired at 270-nm steps with 50–100 ms exposures using an air-cooled QuantEM 512SC CCD camera (Photometrics).

### Image analysis

We quantified endolysosomal luminal pH (ratio of pH-sensitive dextran pHrodo Red 560 to pH-insensitive Dextran-AF640) and ratiometric pulse–chase signals (synuclein–Halo content and aggregate/cytosol ratios) using MATLAB (MathWorks), Fiji (Schindelin et al., 2012), and GPU-accelerated CLIJ2 (Haase et al., 2020). We acquired three-dimensional fluorescence stacks by spinning-disk confocal microscopy and processed them for volumetric segmentation and fluorescence quantification as follows.

#### Background correction

We imported raw 3D stacks into MATLAB and removed background by morphological opening with a spherical structuring element (radius = 10 voxels). We subtracted the resulting background estimate from the raw volume to obtain background-corrected images. *Feature enhancement.* To emphasize punctate and aggregate-like structures, we applied a 3D Laplacian-of-Gaussian filter (fspecial3(’log’,[3 3 2])), followed by Gaussian smoothing (imgaussfilt3, σ = 1). We inverted intensities to highlight compact bright features corresponding to aggregates.

#### Volumetric segmentation

In Fiji/CLIJ2, we segmented objects with the Voronoi–Otsu Labeling algorithm (spot sigma = 0.5; outline sigma = 0.5), which applies local Otsu thresholding to detect bright structures and Voronoi tessellation to separate adjacent objects in 3D. We saved label maps with unique object IDs as 16-bit volumetric TIFFs.

#### Quantification

Using MATLAB, we used the label maps as masks to extract voxel coordinates for each object and measured intensities from the background-corrected images in each fluorescence channel. For every object, we computed integrated fluorescence as the sum of voxel intensities within its mask and calculated channel ratios by dividing the integrated intensity of one channel by that of another for the same object.

#### Data export

For each image, we compiled per-object measurements (object ID, volume, and per-channel integrated fluorescence) and saved the results for downstream analysis.

All segmentation and quantification scripts were written in MATLAB and FIJI are available upon request.

#### pH calibration

We converted the Dextran pHrodo Red 560/Dextran-AF640 ratio to pH using a single-point calibration obtained from cells permeabilized and incubated in pH 7.2 buffer. The measured ratio under these conditions was used to define the region of data with pH of 7.2 or above (neutrality).

#### Cytosol ratiometric analysis

We quantified cytosolic fluorescence ratios with custom MATLAB scripts. For each 3D stack, we extracted the central z-slice and manually drew two rectangular ROIs: (i) a cytosolic ROI avoiding aggregates and (ii) a coverslip-background ROI. For each channel, we computed the background-corrected mean cytosolic intensity as (mean_ROI_cytosol_− mean_RO_Ibackground_. We applied the same ROI coordinates to all channels to ensure spatial registration. We compiled per-channel values across images and calculated the cytosolic ratiometric signal by dividing the background-corrected mean intensity of one channel by that of the other.

Cytosolic fluorescence ratios between the imaging channels were quantified using custom MATLAB scripts. For each 3D image stack, the central z-slice was extracted, and two rectangular regions of interest (ROIs) were manually selected: one within the cytosol (ROI1), avoiding aggregates, and another on the coverslip background (ROI2). The mean fluorescence intensity for each channel was calculated by subtracting the mean intensity of ROI2 from that of ROI1. The same ROI coordinates were automatically applied to all channels to ensure spatial consistency. The resulting per-channel cytosolic differences were compiled across all images for subsequent ratio analysis and the cytosolic ratiometric value was obtained by dividing the mean fluorescence intensity of one channel by that of another.

#### Molecule counting (pulse–chase)

For synuclein–Halo aggregates, we converted integrated fluorescence per object to molecule number by dividing by the single-molecule fluorescence integral. We estimated this single-molecule integral by extrapolating the amplitude from 2D Gaussian fits to Halo–Nup133 spots (from SUM159 cells expressing Halo–Nup133, as described in the NUP study) tracked during photobleaching, yielding the intensity of one fluorophore. We then computed molecular ratios by dividing molecule counts between fluorescence channels for the same object and corrected final counts for an 80% Halo covalent- labeling efficiency.

### FIB-SEM Isotropic Imaging

High-pressure freezing, freeze-substitution, and resin embedding of iNs followed our published protocols ^9,23,37^. Briefly, cells on 6 x 0.1-mm sapphire disks were high-pressure frozen (Wohlwend HPF Compact 03), transferred under liquid nitrogen into osmium tetroxide / acetone substitution medium, processed on an AFS2 warm-up schedule with standard acetone/propylene-oxide to resin exchanges, and polymerized in Embed 812.

The polymerized resin blocks were then mounted on aluminum pin stubs with conductive silver epoxy, platinum-coated (10 nm), and loaded into a Zeiss Crossbeam 540. After eucentric correction and 54° tilt, we deposited a platinum cap, milled a coarse trench (30 kV/30 nA), polished the block face (30 kV/7 nA), and acquired interlaced milling/imaging at 5-nm steps (30 kV/3 nA; SEM 1.5 kV/400 pA) to obtain 5-nm isotropic voxels. Images were from Inlens and ESB detectors (10–15 µs dwell).

The iN sample stained poorly; even a 10× increase in per-voxel beam current during FIB-SEM yielded low SNR. We therefore applied FastFIB ^37^ to improve interpretability and generate 3D volumes suitable for visualization and analysis. Briefly, FastFIB is a self-supervised denoiser based on Noise2Noise ^38^, implemented as a 3D U-Net (four levels; base width 32; kernel size 3; ∼10 M parameters). The blurred network output is compared to a paired noisy target with a Charbonnier loss (α = 0.5; ε = 1×10⁻³), with an added gradient penalty between raw and blurred outputs to suppress high-frequency artifacts. Training used 5×5×5 nm FIB-SEM volumes with intrinsically aligned in-lens secondary and backscattered detector pairs; a brief finetune on the experimental dataset further improved performance.

To aid in the identification and subcellular location of fibrils, we used our ASEM segmentation model ^23^, a supervised 3D algorithm for FIB-SEM volumes. We manually annotated fibrils within a 650 × 270 × 259 voxel ROI (XYZ) with LabKit ^22^ and used these labels to train ASEM for 200,000 iterations (∼23 h on a single A100 GPU, ∼6 GB memory), then generated predictions on the full volumes. The model produced few false negatives but many false positives; we used the positives to locate candidate fibrils to accelerate and facilitate the identification and analysis of fibril-containing regions.

### Statistical Analysis

Although measurements include many cells, the experimental unit was the biological replicate (n = 2). Cells within a replicate are technical measurements, not independent replicates. Accordingly, we report descriptive statistics and replicate-level summaries (plotting all points and showing replicate means where indicated) and did not perform formal inferential testing.

## Data availability

The datasets of raw and denoised FIB-SEM cell images are publicly available as data sets 217.1 and 217.2 at our AWS ASEM data set site http://asem-viewer-env.eba-rrnvmfwa.us-east-1.elasticbeanstalk.com/. This site includes access to the online tool Neuroglancer created to view 3D volumes relevant to this study and others that allows users to quickly select and visualize cells and volumes in the browser.

---

**Video 1 (related to Fig. 3).** 3D FIB-SEM of an endolysosome in an iN prepared by high-pressure freeze substitution, containing a cluster of fibrils. The iN was incubated for one day with 2 µg/ml tau PFF before imaging. A volumetric rendering shows the segmented fibrils (red).

## ACKNOWLEDGMENTS

We thank S.C. Harrison for editorial help and members of the Kirchhausen laboratory for help and encouragement. The research was supported by a National Institute of General Medical Sciences Maximizing Investigators’ Research Award GM130386; by a Massachusetts Life Sciences Pilot grant award to T. Kirchhausen; by the NNF Center of Optimized Oligo Escape and Control of Disease awarded to T. Kirchhausen and N. Hatzakis; by generous support from the Freedom Together Foundation and the Rainwater Foundation to Bradley T. Hayman; by the Cure Alzheimer’s Fund to Bradley T. Hartman and John R. Dickson, by a K08 NIH grant K08AG078341 to John R. Dickson.

Acquisition of the FIB-SEM microscope was supported by a generous grant from Biogen to T. Kirchhausen, and the high-pressure freeze substitution device was made available by S.C. Harrison. Acquisition of the computing hardware including the DGX’s GPU-based computers, CPU clusters, fast access memory, archival servers, and workstations that made possible this study were supported by generous grants from the Massachusetts Life Sciences Center to T. Kirchhausen and by an equipment supplement to the National Institute of General Medical Sciences Maximizing Investigators’ Research Award GM130386. Construction of the server room housing the computing hardware was made possible with generous support from the PCMM Program at Boston Children’s Hospital.

**The authors declare no competing financial interests.**

## Author contributions

Anwesha Sanyal and Tom Kirchhausen conceptualized and designed the experiments; Anwesha Sanyal carried the cell biological experiments; Gustavo Scanavachi carried the optical image analysis; Elliott Somerville and Anwesha Sanyal prepared the samples for FIB-SEM; Elliott Somerville maintained the FIB-SEM and collected and processed the FIB-SEM volumetric data; Jose Da Costa Filho developed and used FastFib, the online tool for 3D data denoising and supported the publicly accessible AWS data repository; Forest Brooks, John R. Dickson, and Bradley T. Hyman generated and characterized the recombinant tau fibrils and human AD brain derived oligomers and fibrils used for the seeding experiments. Tom Kirchhausen drafted the manuscript and finalized it in close consultation with all co-authors.

## Notes

http://asem-viewer-env.eba-rrnvmfwa.us-east-1.elasticbeanstalk.com/

### Competing Interest Statement

The authors have declared no competing interest.

